# SeATAC: a tool for exploring the chromatin landscape and the role of pioneer factors

**DOI:** 10.1101/2022.04.25.489439

**Authors:** Nikita Dsouza, Wuming Gong, Daniel J. Garry

**Affiliations:** Cardiovascular Division, Department of Medicine, University of Minnesota, Minneapolis, MN 55455; Stem Cell Institute, University of Minnesota, Minneapolis, MN 55455; Paul and Sheila Wellstone Muscular Dystrophy Center, University of Minnesota, Minneapolis, MN 55455

## Abstract

The position of the nucleosome and chromatin packaging in eukaryotic genomes govern gene regulation and cellular functions. Assay for Transposase-Accessible Chromatin using sequencing (ATAC-seq) is an efficient and precise method for revealing chromatin accessibility across the genome. However, there is no method that is specifically designed for detecting differential chromatin accessibility using ATAC-seq datasets. In this study, we developed a bioinformatics tool called SeATAC, that used a conditional variational autoencoder (CVAE) model to learn the latent representation of ATAC-seq V-plots, and to estimate the statistically differential chromatin accessibility. We demonstrated that SeATAC outperformed MACS2 and NucleoATAC on four separate tasks including: (1) detection of differential V-plots; (2) definition of nucleosome positions; (3) detection of nucleosome changes and (4) designation of transcriptional factor binding sites (TFBS) with differential chromatin accessibility. By applying SeATAC to several pioneer factor induced differentiation or reprogramming ATAC-seq datasets, we found that induction of these pioneer factors not only relaxed the closed chromatin but also decreased the chromatin accessibility of 20% - 30% of their target sites. These two groups of TF binding sites were characterized by different genomic distribution and histone marks. Here, we present SeATAC as a novel tool to accurately reveal the genomic regions with differential chromatin accessibility from ATAC-seq data.

## Introduction

Eukaryotic genomes are packed into nucleoprotein called chromatin whose basic unit is the nucleosome, which comprises a histone octamer wrapped around 147 base pairs of DNA^1^. Nucleosomes are arranged into regularly spaced arrays, separated by unwrapped linker DNA whose length varies among species and cell types^2^. The dense nucleosome regions (nucleosome occupied regions, NOR) are tightly packed, whereas the loose nucleosome regions (nucleosome free regions, NFR) are more accessible to transcription factors. It is known that precise location of a nucleosome relative to transcriptional target sites can significantly influence factor binding^3–7^. Thus, the chromatin accessibility plays a critical role in regulating gene expression pattern.

High-throughput sequencing techniques such as MNase-seq^8,9^, chemical mapping^10^, DNase-seq^11^, FAIRE-seq^12^ and ATAC-seq^13^ have been developed to assess genome-wide chromatin structure. MNase-seq uses an endo-exonuclease that degrades the accessible linker DNA between nucleosomes and reveals the position of nucleosomes by sequencing the unprotected DNAs. The chemical cleavage method introduces a cysteine substitution at serine 47 in histone H4 (H4S47C) to localize free radical mediated cleavage of nucleosome DNA, followed by performing a copper ion-mediated Fenton reaction to cleave nucleosomal DNAs. The cleaved DNA fragments are then sequenced to estimate the position of the center of the nucleosome. DNase-seq digests with DNase I endonuclease and the resulting DNA fragments correspond to open chromatin region. FAIRE-seq uses formaldehyde and phenol-chloroform extraction separation to isolate nucleosome-depleted DNA from chromatin. Assay for Transposase-Accessible Chromatin using sequencing (ATAC-seq), which utilizes Tn5 transposases to digest accessible genomic DNA, is an efficient and precise method for revealing chromatin accessibility across the genome. Compared with other techniques, ATAC-seq requires less input materials and sample processing time^13,14^, and thus becomes a widely adopted tool for profiling chromatin accessibility of both bulk samples and single cells^15^.

The *fragment size profile* of ATAC-seq paired-end reads can be partitioned into reads generated from putative NFR and NOR regions of DNAs, respectively^13^. The reads from the NOR region have clear periodicity of approximately 150- to 200-bp and produced detailed information on nucleosome position and degree of chromatin compaction^13^. This unique feature of ATAC-seq reads have been utilized to infer the nucleosome positions using NucleoATAC^16^ and deNOPA^17^, which have demonstrated improved performance compared to generic nucleosome calling tools such as DANPOS^18^ and NPS^19^. However, to date, there is no published method for the detection of differential chromatin accessibility specifically for ATAC-seq data. Currently, MACS2^20,21^, which was originally designed for ChIP-seq data, remains the gold standard for analyzing ATAC-seq data^22^, and does not consider ATAC-seq-specific properties.

In this study, we engineered a tool, named SeATAC, to estimate the genomic regions with statistically differential chromatin accessibility from multiple ATAC-seq data. Using SeATAC, each genomic region is represented as a V-plot, a dot-plot showing how sequencing reads with different fragment sizes distribute surrounding one or a set of genomic region(s)^23^. The V-plot based analysis has been used to study nucleosome dynamics flanking the transcription factor (TF) binding sites^23,24^, nucleosome phasing near pioneer factors during reprogramming^25^, clustering the nucleosome profiles near promoters^26^, and examining the distance between nearby nucleosomes^27,28^. However, the V-plot was derived from and visualized for a set of genomic regions due to the noisy and sparse nature of the sequence reads on genomic regions. The difference of V-plots on individual genomic regions between multiple ATAC-seq datasets have never been evaluated before. For SeATAC, we used a conditional variational autoencoder (CVAE) model to learn the latent representation of the ATAC-seq V-plot ^29–31^. With the probabilistic representation of the data, we developed a Bayesian method to evaluate the statistical difference between multiple V-plots. We demonstrated that SeATAC had significantly better performance on four separate tasks compared to MACS2 and/or NucleoATAC on both synthetic and real ATAC-seq datasets. SeATAC is available at https://github.com/gongx030/seatac as an R package.

## Results

### The SeATAC model

The SeATAC model uses a V-plot with a width of 640 bp genomic region and a height of 640 bp of fragment sizes that covers nucleosome free reads (<100 bp), mono-nucleosome reads (between 180 and 247 bp), di-nucleosome reads (between 315 and 473 bp) and tri-nucleosomes (between 558 and 615 bp)^13^. The four groups of ATAC-seq reads represent the majority of total ATAC-seq reads (>95%) and have been successfully used to segment the genomic structure^13,32^. To reduce the impact of noise, an array of 5 × 10 pixels were aggregated together and became a single larger pixel, resulting in an image composed of 128 × 64 pixels. We named the bins along the genomic region dimension and fragment size dimension as *genomic bins* and *fragment size bins*, respectively. The aggregated reads along the genomic bins were then normalized to a vector that sum to one (Figure 1a).

**Figure 1.**
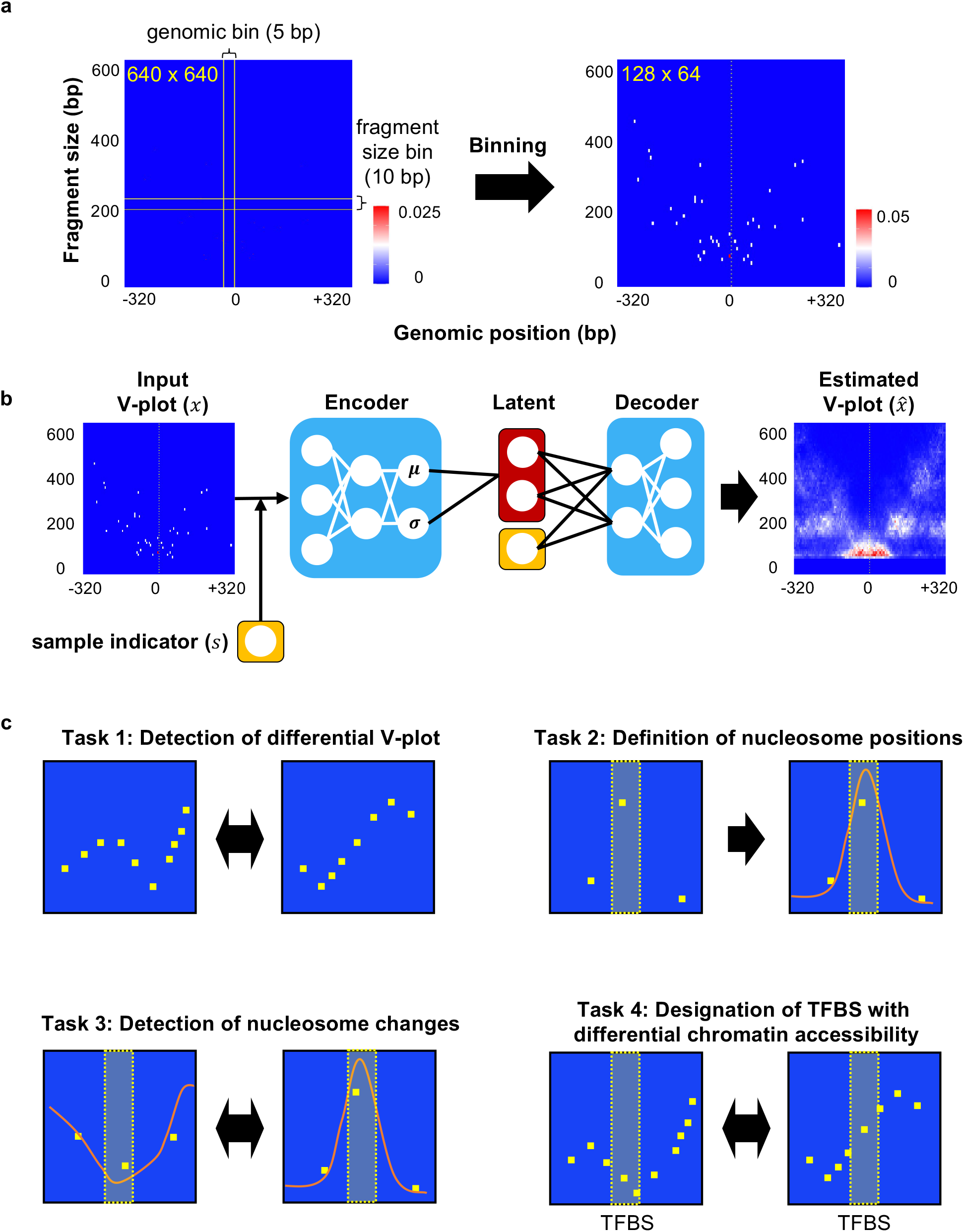
The SeATAC model and tasks for performance evaluation. **(a)** A full V-plot has a width of 640 bp genomic region and a height of 640 bp of fragment sizes (left panel). An array of 5 × 10 pixels are aggregated together and become a single larger pixel, resulting in a 128 × 64 pixels image (right panel). The heatmap color indicates the normalized read density. **(b)** SeATAC models the ATAC-seq V-plot using a conditional variational autoencoder (CVAE) framework. **(c)** Four separate tasks for evaluating the performance of detecting chromatin accessibility changes. MACS2 was excluded from tasks #2 and #3.

We modeled the V-plot ***x**_ni_* of each genomic region *i* in each sample *n* as a probabilistic distribution ***p***(*x_ni_*|***z**_ni_,s_n_*) conditioned on the sample indicator *s_n_* of each sample, as well as an unobserved latent variable ***z**_ni_* (Figure 1b). The sample indicator *s_n_* represents the nuisance variation due to the sample-specific fragment size profile. The latent variable ***z**_ni_* is a *K* dimensional vector of Gaussians representing the remaining variation with respect to the underlying V-plot (*K* = 5). In SeATAC, a neural network serves as a decoder to map the latent variables *z_ni_* and sample indicator *s_n_* to an estimated output V-plot. We expected that latent variables provide batch-corrected representations of the V-plot for the differential analysis. We derived an approximation of the posterior distribution of the latent variable *q*(***z**_ni_*|***x**_ni_,s_n_*) by training another encoder neural network using variational inference and a scalable stochastic optimization procedure^29,30^. The variational distribution *q*(***z**_ni_*|***x**_ni_,s_n_*) is chosen to be Gaussian with a diagonal covariance matrix, where the mean and covariance are estimated by an encoder neural network applied to (***x**_ni_,s_n_*). The variational evidence lower bound (ELBO) is

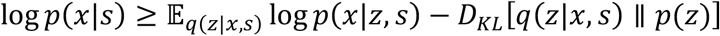

A standard multivariable normal prior *p*(***z**_ni_*) is used in SeATAC because it can be reparametrized into a way that allows backpropagation to flow through the deterministic nodes^29^. To optimize this lower bound we used the reparameterization trick to compute low-variance Monte Carlo estimates of the expectations’ gradients. Throughout the study, we used Adam optimizer (learning rate = 0.01) with a cosine learning rate schedular with warmup.

### SeATAC corrects batch effects in ATAC-seq data

Although the fragment size profile (the fragment size density plot) provided similar fragment length estimation regrading NFR and nucleosomes (mono-nucleosomes, di-nucleosomes, tri-nucleosomes, etc.)^13^, the exact pattern differed across ATAC-seq datasets, resulting in different fragment size ranges and density for NFR and nucleosome reads. We assumed that the majority of the batch effects in the ATAC-seq were due to the difference of the fragment size profile^13^. In the SeATAC model, an embedding layer first maps the sample indicator *s_n_* to the fragment size vector *g_n_* and combines with the input V-plot to produce a modified V-plot. Then, convolutional neural networks (CNN) maps the modified V-plot to the latent variables. Once the model was optimized, SeATAC used a constant sample indicator *s*_0_ to replace the sample specific indicator *s_n_* to generate a batch-free estimated V-plot.

We applied SeATAC to a human hematopoietic differentiation datasets with 13 samples^33^, and each sample showed a distinct fragment size profile (Supplementary Figure 1a). We randomly sampled 2,000 640-bp genomic regions, generated batch-free V-plot and computed the aggregated fragment size profile by averaging along each fragment size bin. The corrected fragment size profile became consistent across 13 samples, suggesting that SeATAC was able to successfully correct the batch effects due to difference in fragment size profile, allowing SeATAC, in an unbiased fashion, to compare multiple ATAC-seq samples.

### Tasks for performance evaluation

We designed four separate tasks for evaluating the performance of detecting chromatin accessibility changes including: (1) the detection of differential V-plots; (2) the recovery of nucleosome positions from sparse ATAC-seq data; (3) calling differential nucleosomes and (4) the designation of transcriptional factor binding sites (TFBS) following increased chromatin accessibility (Figure 1c). The task #1 was to determine whether or not the V-plot for a genomic region was different between multiple ATAC-seq samples. The task #2 and #3 asked the methods to recover (task #2) and to compare (task #3) nucleosome positions. We excluded MACS2 from these two tasks since MACS2 was not capable of calling the nucleosomes directly. Tasks #1-#3 were evaluated on the datasets down-sampled from a full ATAC-seq dataset. Task #4 focused on the detection of individual TFBS with differential chromatin accessibility and was evaluated using several ATAC-seq datasets of TF induced reprogramming.

### SeATAC detects differential V-plot

To define a benchmark dataset for testing a differential V-plot, we generated two separate down-sampled datasets (dataset #1 and dataset #2) that included 10% of sequencing reads of a full ATAC-seq dataset (GM12878) by using different random seeds, separately. Then every read in dataset #2 was shifted to 3′ direction by a pre-specified distance (e.g. 100bp) to generate a new dataset #3. Thus, dataset #1 and dataset #2 should have the identical V-plot for any genomic regions, while dataset #1 and dataset #3 should have different V-plot because the shift size is smaller than the length of nucleosome DNAs and the linker DNAs (Figure 2a). We used SeATAC, MACS2 and NucleoATAC to compare dataset #1 vs. dataset #2, and dataset #1 vs. dataset #3, and evaluate the performance of calling differential V-plots by computing the receiver operating characteristic (ROC) curves, respectively (Figure 2b). The SeATAC *p*-values (*p^SeATAC^*), MACS2 *p*-value (*p^MACS2^*) and the maximum difference of NucleoATAC signal were used to rank the differential V-plots (see Methods). We generated six different pairs of down-sampled datasets (#1 - #3), and evaluated the performance on different shift size for dataset #3 (10, 20, 50 and 100 bp). The average Area Under the ROC Curve (AUC) of SeATAC was 64.8, 67.2, 70.7 and 79.5 for shift size of 10, 20, 50 and 100 bp, respectively. In comparison, the AUC of MACS2 was below 50, while the AUC of NucleoATAC ranged from 50.0 to 54.3 (Figure 2c). These results suggested that SeATAC was able to detect the differential V-plot across different shift sizes.

**Figure 2.**
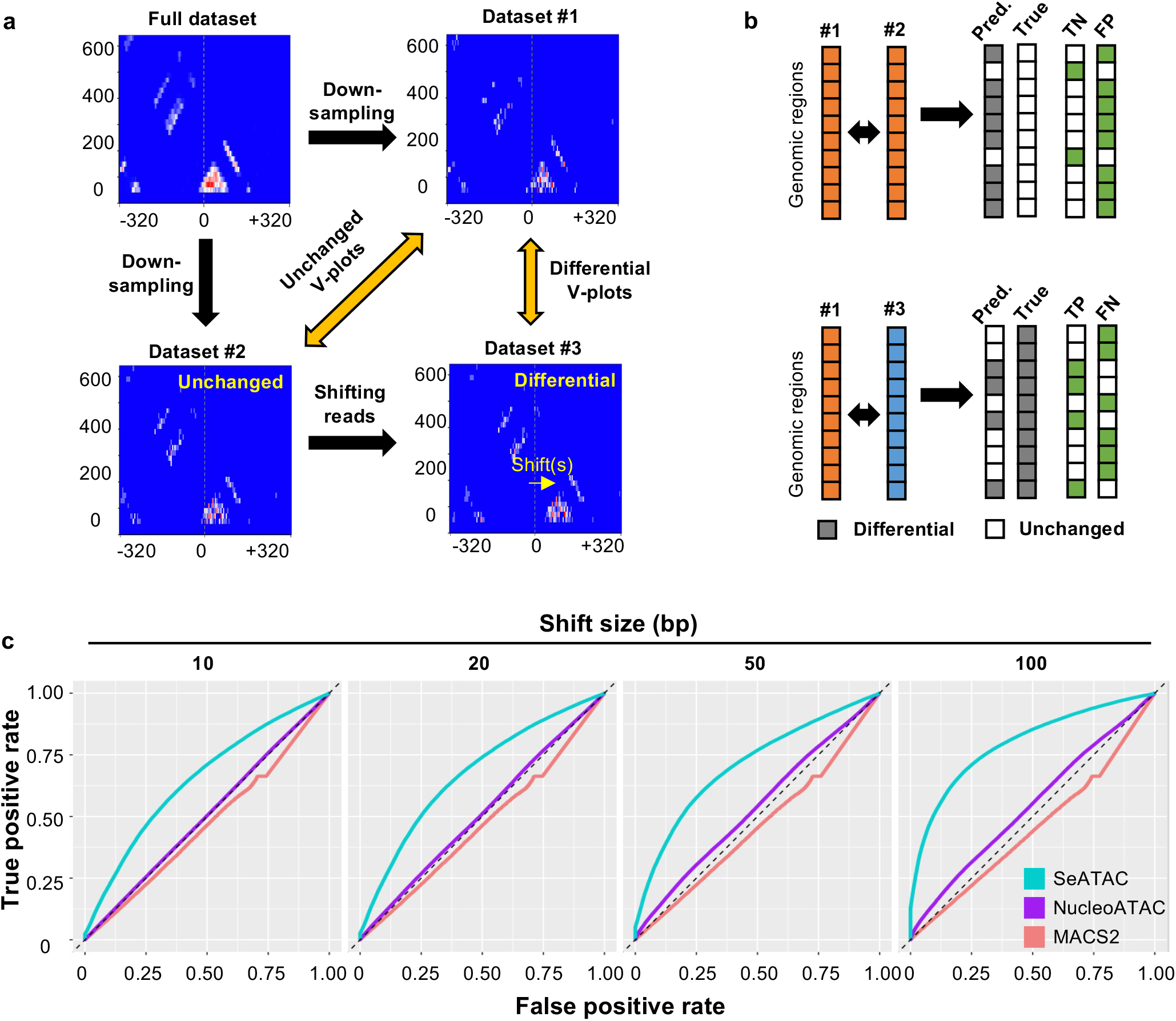
SeATAC detects differential V-plots. **(a)** A full ATAC-seq dataset is down-sampled to two separate datasets (dataset #1 and dataset #2) that includes 10% of the sequencing reads. Every read in dataset #2 is shifted to the 3′ direction by a pre-specified distance (e.g. 100bp) to generate a new dataset #3. The dataset #1 and dataset #2 have the identical V-plot for any genomic regions, while dataset #1 and dataset #3 have different V-plots. **(b)** Different tools are used to compare dataset #1 vs. dataset #2, and dataset #1 vs. dataset #3 to detect differential V-plots. The true positive (TP), false positive (FP), true negative (TN) and false negative (FN) predictions are computed. The receiver operating characteristic (ROC) curve is used to evaluate the performance of different tools. **(c)** The ROC curves for SeATAC, NucleoATAC and MACS2 for four different shift sizes: 10, 20, 50 and 100 bp are shown.

### SeATAC recovers nucleosome positions from sparse ATAC-seq data

To evaluate how well SeATAC detected nucleosome positions from sparse ATAC-seq data, we first defined the NFR or NOR positions on a full ATAC-seq dataset (GM12878). A genomic locus was considered as a NOR center if the NucleoATAC signal at this locus was greater than 0.5 and was also greater than any other positions in the flanking 200 bp region. A genomic locus was considered as a NFR center if the NucleoATAC signal at this locus was smaller than 0.01 and was also smaller than any other position in the flanking 200 bp region. There were 9,965 and 316,075 NOR and NFR centers in the full ATAC-seq data. We randomly sampled ~5,000 NOR and NFR centers to evaluate the performance of nucleosome calling. We down-sampled the full ATAC-seq dataset to 0.1%, 1% and 10% of the full datasets and used SeATAC and NucleoATAC to estimate the nucleosome signals at each NOR and NFR centers (see Methods). SeATAC demonstrated overall superior performance on calling nucleosomes from sparse ATAC-seq data with AUR of 0.583, 0.606 and 0.653 for 0.1%, 1% and 10% down-sampled datasets, respectively, while the AUC for NucleoATAC were 0.503, 0.491 and 0.591, respectively (Figure 3a). Among ~5,000 NORs, we identified 2,042, 3,453 and 22 regions that were called by both SeATAC and NucleoATAC, SeATAC only and NucleoATAC only as nucleosomes, respectively (*Nuc^SeATAC^* > 0.5 or *Nuc^NucATAC^* > 0.2). The center of these genomic regions that were called as nucleosomes by SeATAC only showed enriched nucleosome signals supported by both NucleoATAC estimation on the full dataset and an MNase-seq dataset on GM12878^34^ (Figure 3b). These results supported the notion that SeATAC had better performance on estimating nucleosomes from sparse ATAC-seq data.

**Figure 3.**
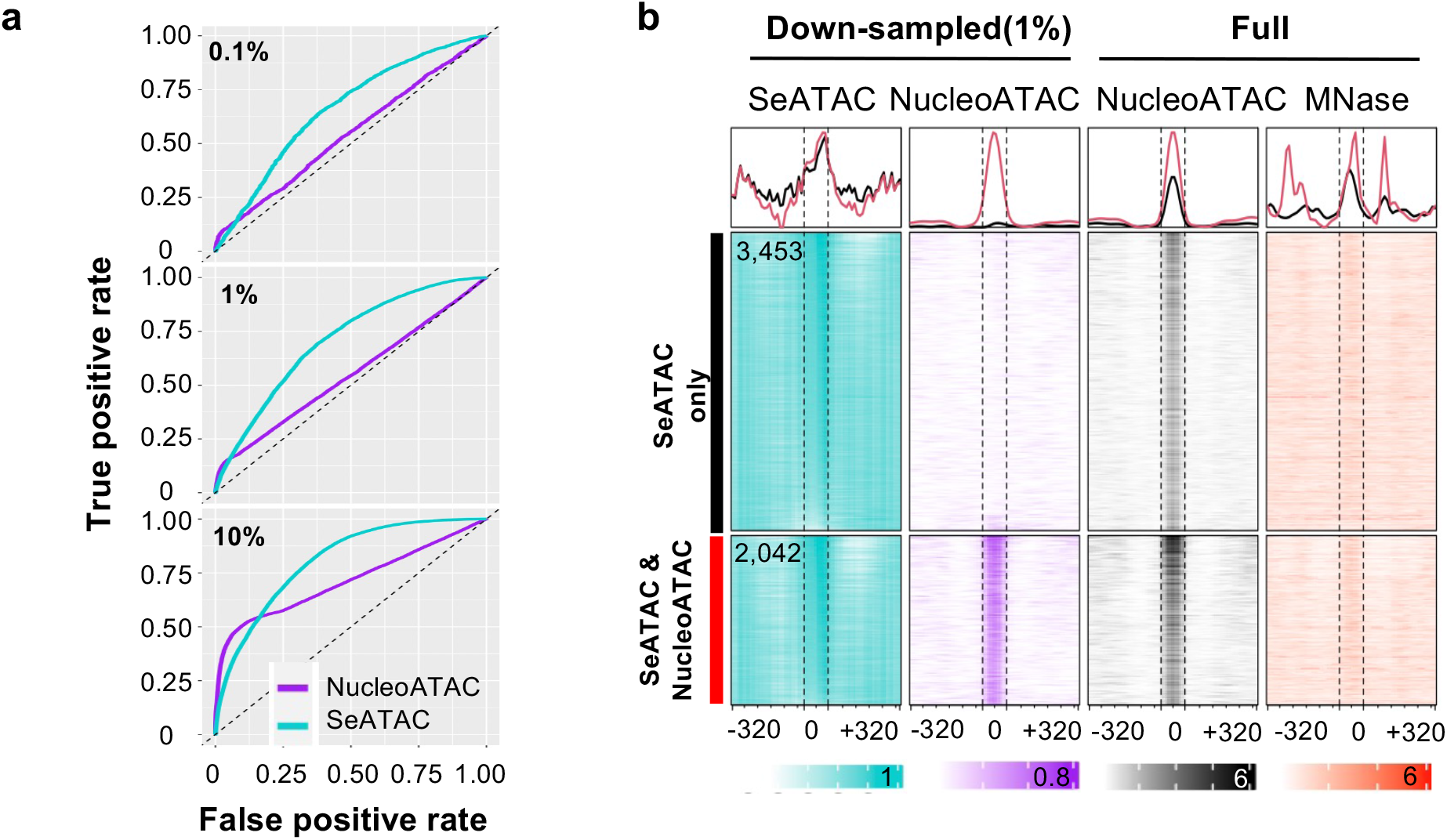
SeATAC recovers nucleosome positions from sparse ATAC-seq. **(a)** The ROC curve for recovering nucleosome positions from ATAC-seq with 0.1%, 1% and 10% of the sequencing reads randomly sampled from the full dataset (GM12878). **(b)** The heatmaps shows the nucleosome density estimated by SeATAC (blue) and NucleoATAC (purple) on a 1% down-sampled dataset. There are 2,042 and 3,453 regions (640 bp) identified by both SeATAC / NucleoATAC, and by SeATAC only as nucleosomes. The NucleoATAC signal on the full dataset (black) and a MNase-seq dataset on GM12878 (red) for these regions are also shown.

### SeATAC detects nucleosome changes

By using the ground truth NOR and NFR centers on full GM12878 ATAC-seq dataset, we could also evaluate how capable SeATAC was regarding the call of the nucleosome change from NFR to NOR. We randomly sampled 5,000 NFR/NOR pairs and applied SeATAC and NucleoATAC to evaluate the nucleosome changes at the center of each NFR/NOR pairs on a down-sampled ATAC-seq dataset with 10% of sequencing reads. The nucleosome changes were ranked by SeATAC’s *differential central nucleosome score* (δ^*SeATAC*^) and NucleoATAC’s *differential central signal* (δ^*NucATAC*^), respectively (see Methods). SeATAC demonstrated superior performance on calling nucleosome changes than NucleoATAC with an AUC of 0.904 vs. 0.827 (Figure 4a). Among ~ 5k NFR/NOR pairs, SeATAC was able to successfully identify more than 72.9% of genuine NFR/NOR changes compared to NucleoATAC (1,278 vs. 739), and these changes were supported by the NucleoATAC signals on the full dataset and an MNase-seq dataset ^34^ (Figure 4b and 4c). These results suggested that SeATAC could more accurately detect the nucleosome changes between ATAC-seq samples.

**Figure 4.**
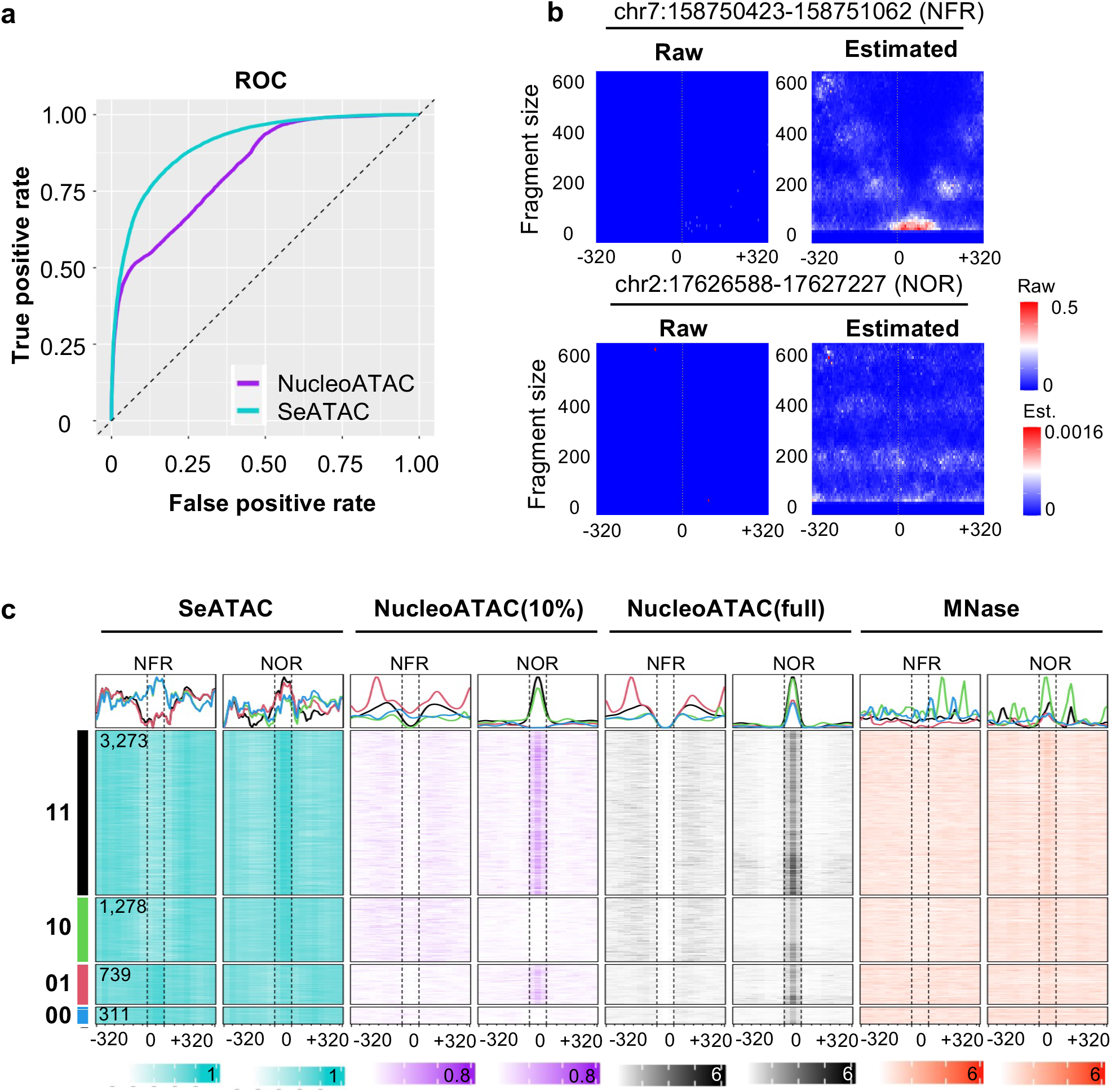
SeATAC detects nucleosome changes. **(a)** The ROC curve for detecting nucleosome changes from ATAC-seq with 10% of the sequencing reads from the full dataset (GM12878). **(b)** The raw and estimated V-plot of a NFR (chr1:113162059-113162698) and a NOR (chr2:226653061-226653700) region are shown. The heatmap color indicates the normalized read density. **(c)** The heatmaps show the nucleosome density of ~5,000 sampled NOR and NFR regions estimated by SeATAC (blue) and NucleoATAC (purple) on a 10% down-sampled dataset. There are 3,276, 1,278, 739 and 311 regions that are identified as a change from NFR to NOR (with decreased chromatin accessibility) by both SeATAC and NucleoATAC (11), by SeATAC only (10), by NucleoATAC only (01) and by neither of them (00), respectively. The NucleoATAC signal on the full dataset (black) and a MNase-seq dataset on GM12878 (red) for these regions are also shown.

### SeATAC detects chromatin accessibility changes associated with biological functions

Having established SeATAC’s superior performance on three separate tasks using synthetic data, we then applied SeATAC to ATAC-seq datasets of Etv2 induced MEF reprogramming and ES/EB differentiation^35^. Etv2 is an essential transcription factor for the development of cardiac, endothelial and hematopoietic lineages^36–46^. Moreover, Etv2 has recently been shown to function as a pioneer factor. In these studies, the induction of Etv2 drove embryonic body (EB) and MEFs to an endothelial fate^35^. Therefore, we hypothesized that the relaxed Etv2 binding sites (becoming more accessible) during the Etv2 induced differentiation or reprogramming period and should be closely associated with the endothelial function.

SeATAC identified 5,451 and 2,142 Etv2 motifs with increased chromatin accessibility from MEF reprogramming (undifferentiated MEFs vs. Flk1^+^ cells at 7 days post induction) and EB differentiation (D2.5 EB vs. Flk1^+^ cells at 12 hours post induction) ATAC-seq data, respectively (adjusted *p*-value < 0.05 and *δ^NOR^* < −0.1). Interestingly, SeATAC identified 2,776 and 1,626 relaxed Etv2 motifs that were detected by neither MACS2 nor NucleoATAC The aggregated V-plot of 1,626 SeATAC-only Etv2 binding sites showed increased NFR reads, while the aggregated V-plot of 222 MACS2-only and 2,305 NucleoATAC-only Etv2 binding sites did not show significant changes from undifferentiated EBs to Flk1^+^ cells from 12 hours post induction (Figure 5c). The aggregated V-plot of SeATAC-only, MACS2-only and NucleoATAC-only Etv2 binding sites from MEF reprogramming also showed a similar pattern (Supplementary Figure 2a). Moreover, the pathway analysis showed that the relaxed Etv2 binding sites identified by SeATAC were more significantly associated with Gene Ontology terms related to endothelial development and cell migration (Figure 5d).

**Figure 5.**
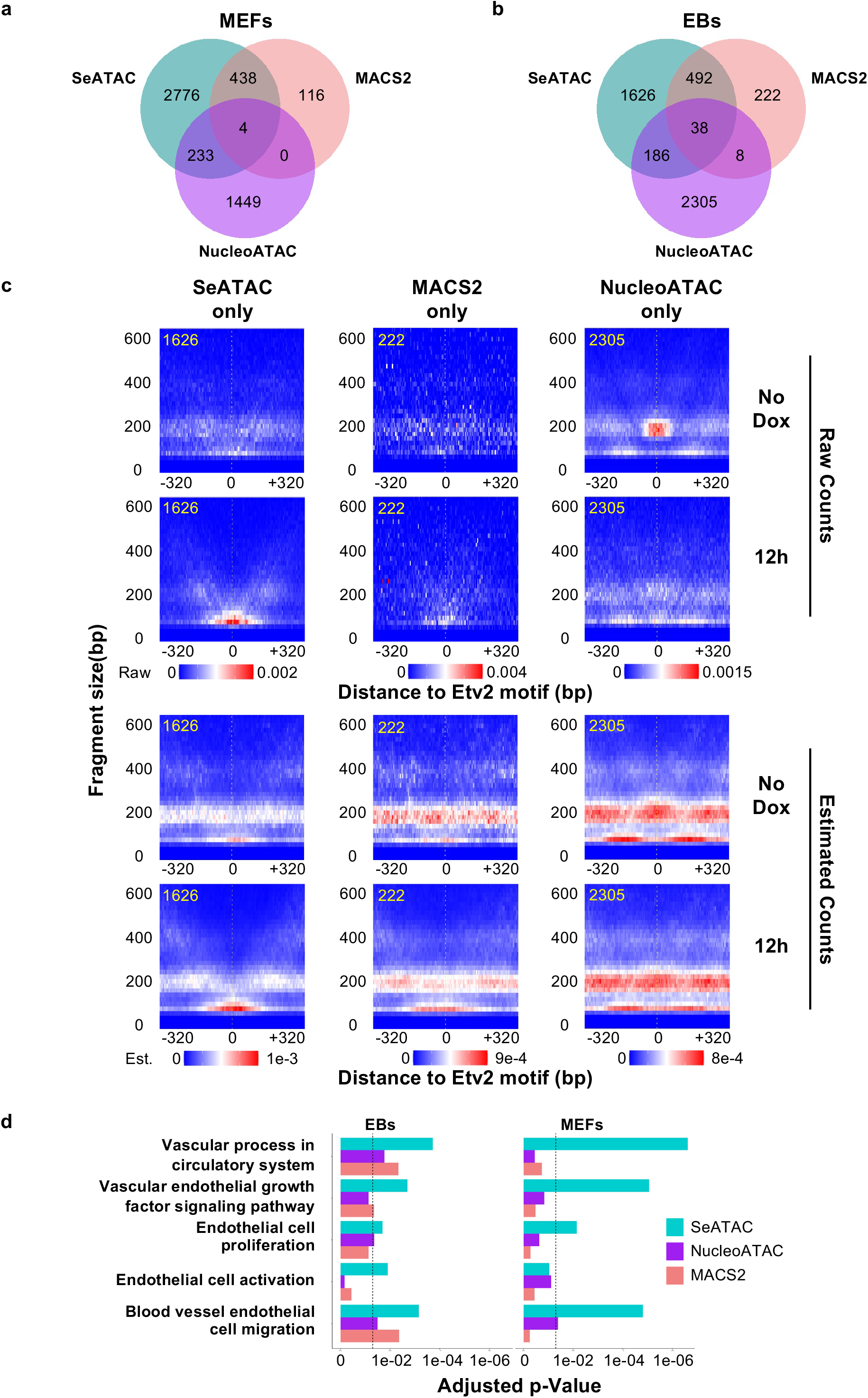
SeATAC detects Etv2 binding sites with increased chromatin accessibility during Etv2 induced EB differentiation and MEF reprogramming. **(a-b)** The Venn diagrams show the number of Etv2 motifs with increased chromatin accessibility identified by SeATAC, MACS2 and NucleoATAC, in **(a)** Etv2 induced MEF reprogramming (undifferentiated MEFs vs. Flk1^+^ cells at 7 days post-induction) and **(b)** Etv2 induced EB differentiation (D2.5 EB vs. Flk1^+^ cells at 12 hours post-induction). **(c)** The aggregated V-plot includes 1,626, 222 and 2,305 Etv2 motifs with increased chromatin accessibility identified by SeATAC only, MACS2 only and NucleoATAC only in ATAC-seq data of Etv2 induced EB differentiation (day 2.5 EB vs. Flk1^+^ cells at 12 hours post-induction). Both raw V-plots and estimated V-plots are shown. The heatmap color indicates the normalized read density for raw counts (top) and the estimated read density for estimated read counts (bottom). **(d)** The barplots show the Gene Ontology (GO) terms that are significantly associated with the genes where the promoters (−5,000 - +1,000bp region flanking the TSS) have Etv2 motifs with increased chromatin accessibility, identified by SeATAC, MACS2 and NucleoATAC. The y-axis showed the adjusted *p*-value of the pathway analysis.

We examined two additional ATAC-seq datasets of Ascl1 induced neural reprogramming^25^ (undifferentiated MEFs vs. 22 days post induction of Ascl1) and OSK (Oct4, Sox2 and Klf4) induced reprogramming^47^. We found that the relaxed Ascl1 binding sites identified by SeATAC were more significantly associated with Gene Ontology terms related to neurogenesis and neuron migration (Supplementary Figure 3), while the relaxed OSK binding sites were more significantly associated with Gene Ontology terms related to stem cell development, fibroblast growth factor receptor signaling pathways and canonical Wnt signaling pathway (Supplementary Figure 4)^47,48^. These results suggested that SeATAC was able to identify TFBS with differential chromatin accessibility and closely related biological functions. Importantly, these differential TFBS were missed by conventional tools such as MACS2 and NucleoATAC.

### Induction of pioneer factors cause both chromatin relaxation and closure

Previous studies showed that pioneer factors such as Etv2, Ascl1 and OSK could recognize their target DNA sequences in compacted chromatin, recruit chromatin remodelers and trigger the relaxation of the adjacent chromatin landscape to accommodate non-pioneer transcription factors^49,50^. In Etv2 induced EB differentiation and MEF reprogramming, although the overall Etv2 motif associated chromatin accessibility significantly increased, as suggested by chromVAR analysis^51^, SeATAC showed that among the Etv2 motifs with differential chromatin accessibility, ~30% (24.6% in MEFs and 35.1% in EBs) the Etv2 motifs showed decreased chromatin accessibility during Etv2 induced differentiation (Figure 6a and Figure 6c). We found that a majority of the Etv2 motifs with decreased chromatin accessibility were located near the promoter regions (Figure 6b), and marked by euchromatic marks such as H3K4me1, H3K4me2, H3K27ac and P300 (Figure 6d). The decrease of chromatin accessibility was also coupled with the decrease of Brg1 (SMARCA4) density, a key SWI/SNF-related chromatin-remodeling complex that facilitates chromatin relaxation (Figure 6d)^52^. Additionally, we found that the genes, which harbor Etv2 binding sites with decreased chromatin accessibility (NFR->NOR) in the promoter regions (−5,000 - +1,000 bp surrounding the TSS), including Brachyury (T) and Mycn, were more likely to be down-regulated during the differentiation process (Figure 6e-6g, Supplementary Figure 2c), suggesting the Etv2 may regulate gene expression by reducing the chromatin accessibility of their binding sites.

**Figure 6.**
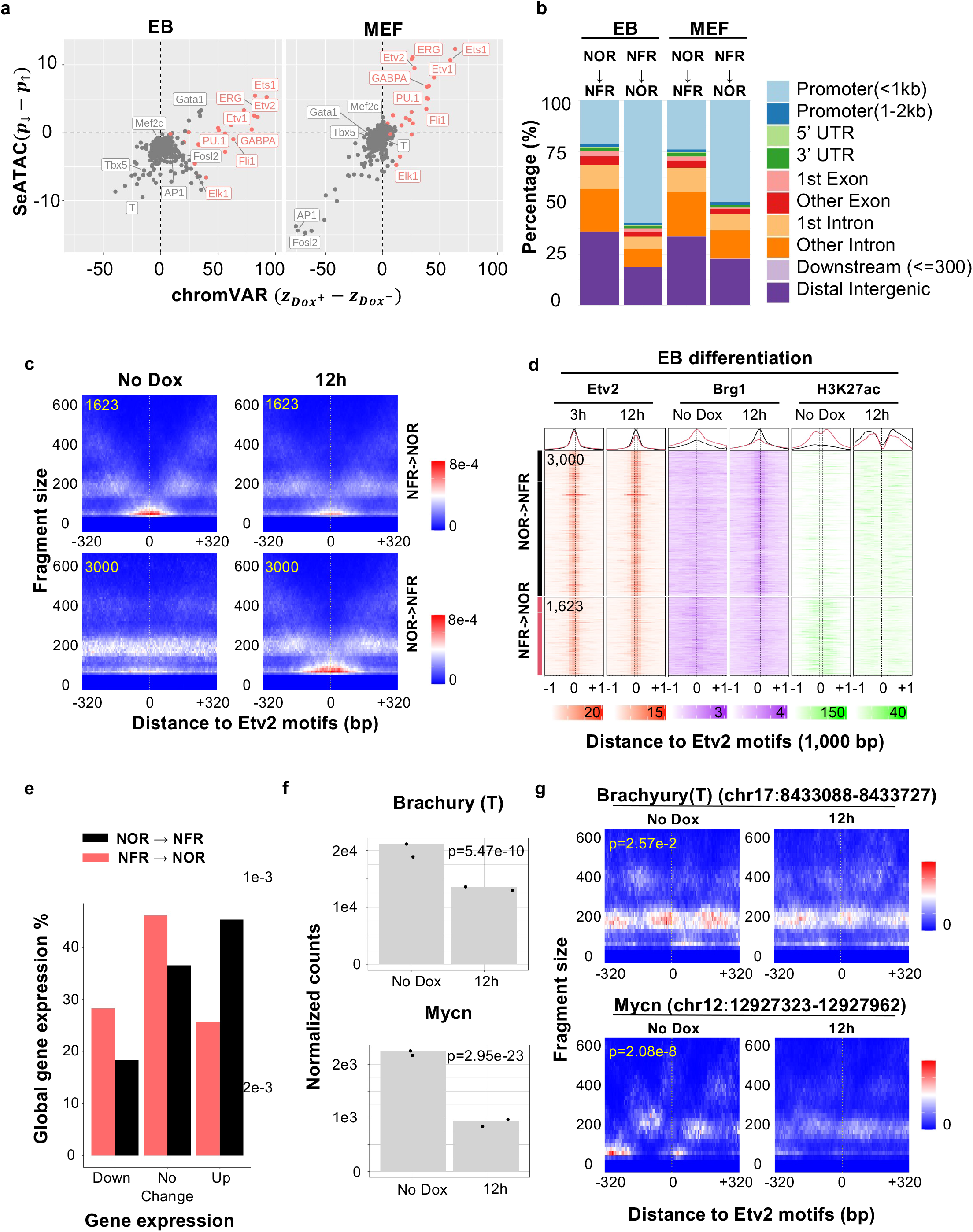
Inducing Etv2 causes both chromatin relaxation and closure at Etv2 binding sites. **(a)** The dot plots compare the changes of motif associated chromatin accessibility estimated by chromVAR (x-axis) and the difference of the percent of TFBS with decreased or increased chromatin accessibility estimated by SeATAC (y-axis). ***z_dox_***+ and ***z_Dox_***− are the normalized deviation score of Dox^+^ condition (Flk1^+^ cells at 7 days post-induction for MEF reprogramming or Flk1^+^ cells at 12 hours post-induction for EB differentiation) and Dox>^−^ condition (undifferentiated MEFs or D2.5 EBs). ***p***_↓_ and ***p***_↑_ are the percent of TFBS that shows decreased or increased chromatin accessibility in Dox^+^ condition compared with the Dox^−^ condition. **(b)** The barplots show the genomic distribution of Etv2 binding sites with decreased (NFR->NOR) or increased (NOR->NFR) chromatin accessibility in EB differentiation or MEF reprogramming. The change of chromatin accessibility is estimated by SeATAC. **(c)** The aggregated V-plot include 3,000 and 1,623 Etv2 binding sites that have increased (NOR->NFR) or decreased (NFR->NOR) chromatin accessibility during MEF reprograming. The heatmap color indicates the estimated read density. **(d)** The heatmaps show the Etv2, Brg1, H3K27ac ChIP-seq of 3,000 and 1,623 Etv2 binding sites that have increased (NOR->NFR) or decreased (NFR->NOR) chromatin accessibility at day 2.5 EB (Brg1 and H3K27ac), 3 hours post Etv2 induction (Etv2), and 12 hours post Etv2 induction (Etv2, Brg1 and H3K27ac). The change of chromatin accessibility is estimated by SeATAC. **(e)** The barplots show the percent of genes that were down-regulated, up-regulated or not changed between day 2.5 EB and 12 hours post Etv2 induction. **(f-g)** Brachyury (T) and Mycn **(f)** are significantly down-regulated during the Etv2 induced differentiation and **(g)** have Etv2 motifs that become significantly less accessible during differentiation at their promoter region (−5,000 - +1,000bp region flanking the TSS). The heatmap color indicates estimated read density.

The analysis of the ATAC-seq dataset of Ascl1 induced neural reprogramming^25^ revealed among Ascl1 motifs with differential chromatin accessibility, 19.8% showed decreased chromatin accessibility (Supplementary Figure 5a-5d). Similar to Etv2 motifs, the Ascl1 motifs with decreased chromatin accessibility (NFR->NOR) were marked by euchromatic histone marks (Supplementary Figure 4b and 4c) and were present in the promoters of genes that were down-regulated during the reprogramming, including Hmga2^53^, Egfr^54^, Elf4^55^, as well as Notch signaling member Hes1^56^ (Supplementary Figure 5e). The analysis of the ATAC-seq dataset of OSK induced reprogramming ^47^ also revealed that the OSK motifs that became less accessible (NFR->NOR) during the reprograming were marked by euchromatic marks in MEFs, more likely located at the promoter regions, and present at the promoters of down-regulated genes during reprogramming, including Maf^57^ and Smad3^58^ (Supplementary Figure 6).

These results clearly showed that pioneer factors could recognize DNA sequences in both closed and open chromatin structure and alter the chromatin landscape in a context dependent manner.

## Discussion

In the present study, we presented a novel algorithm SeATAC for the detection of genomic regions with differential chromatin accessibility and nucleosome positions. SeATAC employed a conditional variational autoencoder framework to model the ATAC-seq-specific V-plot while addressing the batch effect in the ATAC-seq data, allowing an unbiased comparison across multiple samples. The convolutional neural network (CNN) blocks used in the encoder network allowed SeATAC to robustly estimate the posterior distribution of the latent variables by considering ATAC-seq specific fragment size profile, resulting in superior performance on several tasks such as detecting differential V-plot, recovering nucleosome positions, detecting nucleosome changes and calling TFBS with differential chromatin accessibility compared to conventional methods such as MACS2 and NucleoATAC.

When applying ATAC-seq datasets on TF induced differentiation and reprogramming methods, SeATAC more accurately identified TFBS with differential chromatin accessibility, resulting in a more significant association with the underlying biological function. Surprisingly, we found that the induction of pioneer factors such as Etv2, Ascl1, Oct4, Sox2 and Klf4 not only relaxed the compacted chromatin surrounding the respective binding sites, but also resulted in the reduction of chromatin accessibility near 20% ~ 30% of the binding sites. The mechanism of pioneer factor induced chromatin closure and their roles in lineage specification has never been explored before and it warrants further investigation.

The SeATAC framework can be extended to model single cell ATAC-seq (scATAC-seq) data and to investigate the V-plot dynamics in the scATAC-seq data^15^. More sophisticated neural architecture such as attentions or transformer encoders^59^ can be used to replace CNN layers to better model the dependences of ATAC-seq reads on V-plot^60^. Although throughout this study, a default width of 640 bp was used for the V-plot, a wider V-plot (e.g. 2,048 bp) can be potentially used to model more nucleosomes at a specific locus and distant dependencies. We believe that SeATAC provides an accurate and powerful way of revealing chromatin dynamics from the ATAC-seq data and be a valuable tool to examine the chromatin landscape and the functional role of epigenetic regulators.

## Methods

### Neural architecture

For each genomic region *i* in *S* ATAC-seq samples, the V-plot with the dimension of *W* × *H* × 1 from each sample was stacked together at the channel dimension to form an array ***X**_i_* ∈ R^*W* ×*H*×*S*^, where *W* is the number of genomic bins, *H* is the number of fragment size bins, and *S* is the sample size. SeATAC used *W*=128 and *H*=64 by default. An embedding layer first maps the sample indicator ****s**** ∈*Z*^*S*^ to a fragment size array *g* ∈ R1×*H*×*S*. An encoder neural network then maps the modified V-plot (***X**_i_* + ***g***) to latent variables with the mean of ***Z**_i_* ∈ R^*K*×*S*^ and the standard deviation of ***σ*** ∈ R^*K*×1^, where *K* is the dimension of the latent variable (*K* = 5 by default). The encoder network consists of four convolutional neural networks (CNN) blocks, where each block consists of a CNN layer (filter of 16, stride of 2 and kernel size of 3), a batch normalization layer and an Rectified linear Unit (ReLU) activation layer. The output of the CNN blocks are flattened and mapped to latent variables with the mean of ***z**_i_* and standard deviation of ***σ*** by a dense layer. The decoder neural network first maps concatenated latent variable ***z**_i_* and sample indicator ***s*** to a vector of 128 by a dense layer, followed by four transposed CNN blocks, where each block consists of a transposed CNN layer (filter of 16, stride of 2 and kernel size of 3), a batch normalization layer and an Rectified linear Unit (ReLU) activation layer. The output of the CNN blocks feed into a final softmax activation layer to normalize the values in each genomic bin to a vector that sum to one. In this study, we employed the binary cross entropy loss to minimize the difference between input and the estimated V-plot.

### Task #1: Detection of differential V-plots

#### SeATAC

With its probabilistic representation of the data, SeATAC provides a natural way of testing differential V-plot, while intrinsically controlling for nuisance factors. We used the SeATAC model to approximate the posterior probability of the batch-free latent variable ***z***. For each genomic region and a pair of ATAC-seq samples with latent variables of mean of (*z_ak_,z_bk_*) and variance of 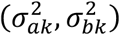, where *k* = 1, … *K* and *K* is the dimension of the latent variables, we constructed a *χ*^2^ variable *Q* by standardizing the difference between ***z**_a_* and ***z**_b_*:

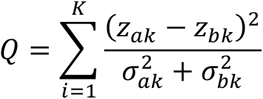

This *χ*^2^ variable *Q* measures the standardized distance between a pair of V-plot on the latent space and a *χ*^2^ test with *K* degree of freedom was used to compute a *p*-value of the difference between two V-plot^61^ (*p^SeATAC^*).

#### MACS2

We used MACS2 (v2.1.1)^20,21^ to compare two BAM files (file1.bam and file2.bam) twice by swapping the control and treatment samples, using the following parameters: “macs2 callpeak -q 0.05 --call-summits -f BAMPE --nomodel -t file1.bam -c file2.bam --keep-dup all” and “macs2 callpeak -q 0.05 --call-summits -f BAMPE -nomodel -t file2.bam -c file1.bam --keep-dup all”. The results of two runs were merged and the minimum *p*-values of all summits that overlapped with a 640-bp genomic region was used as *p*-values for this genomic region (*p^MACS2^*).

#### NucleoATAC

We used NucleoATAC (v0.3.4 with default parameters)^16^ to estimate the nucleosome signal of two BAM files separately, and calculated the difference of estimated nucleosome signal for genomic regions. The maximum absolute values of the difference of nucleosome signals that overlapped with a 640-bp genomic region was used as the difference of nucleosome signals for this genomic region.

### Task #2: Estimating the nucleosome signals

#### SeATAC

For any genomic region, SeATAC generates estimated V-plot 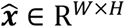 based on the latent variables ***z*** and a constant sample indicator *s*_0_, from which we computed the *central NFR score*:

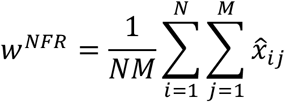

where *N* is the number of central genomic bins and *M* is the number of fragment size bins for NFR. The central genomic bins were defined as the genomic bins which distance to the V-plot center (*d_i_*) is less than 50 bp (−50 ≤ *d_i_* ≤ 50), and fragment size bins for NFR (*f_j_*) were defined as the fragment size less than 150 bp (*f_j_* ≤ 150). The *center nucleosome score* were defined as:

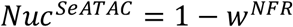

The central nucleosome score (*Nuc^SeATAC^*) was used as the nucleosome score estimated by SeATAC to rank the nucleosomes.

#### NucleoATAC

We used NucleoATAC to estimate the nucleosome signal from the input BAM files. We defined the *central NucleoATAC signal* as the average NucleoATAC signal over 100bp region flanking the V-plot center.

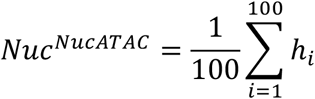

where *h_i_* is the NucleoATAC signal at position *i*. We used central NucleoATAC signal (*Nuc^NucATAC^*) to rank the nucleosomes for this task.

### Task #3: Detection of nucleosome changes

#### SeATAC

For any genomic region between a pair of ATAC-seq samples (*i,j*), SeATAC computed the *differential central nucleosome score* by:

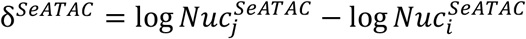

δ^*SeATAC*^ quantitatively measures how estimated nucleosome signal changes from sample *i* to *i* over the 100 bp regions flanking the center.

#### NucleoATAC

For any genomic region between a pair of ATAC-seq samples (*i,j*), the *differential central NucleoATAC signal* (δ^*NucATAC*^) was defined as the difference of the average NucleoATAC signal over 100bp region flanking the V-plot center between a pair of ATAC-seq samples:

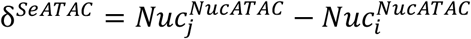

### Task #4: Designation of TFBS with increased chromatin accessibility

#### SeATAC

Between a pair of ATAC-seq samples (*i,j*), SeATAC determined that a TFBS became more accessible in sample *j* compared with sample *i* if the adjusted p-value < 0.05 and δ^*SeATAC*^ < −0.1.

#### MACS2

First, we used MACS2 to compare the sample *i* (file1 .bam) and sample *j* (file2.bam) using the following parameters: “macs2 callpeak -q 0.05 --call-summits -f BAMPE --nomodel -c file1.bam -t file2.bam --keep-dup all”. Then we computed MACS2 *p*-value for a specific TFBS as the minimum *p*-values of all summits that overlapped with 100 bp region flanking this TFBS (*p^MACS2^*). MACS2 determined that a TFBS became more accessible if adjusted *p^MACS2^* < 0.05.

#### NucleoATAC

NucleoATAC determined that a TFBS became more accessible in sample *j* compared with sample *i* if the δ^*NucATAC*^ < −0.4.

### Data availability

#### Human hematopoietic differentiation

The dataset was downloaded from NCBI GEO database (GSE96771). A union set with 491,437 peaks defined by the original authors were used for the downstream analysis.

#### GM12878

The human EBV-transformed lymphoblastoid cell line (LCL) ATAC-seq data were downloaded from NCBI GEO database (GSE47753). The sequence reads from three replicates of 50k cell sample (GSM1155957, GSM1155958 and GSM1155959) were pooled and used for the downstream analysis. The 86,004 peaks called by MACS2(v2.1.1)^20,21^ were used for the downstream analysis.

#### Etv2 induced MEF reprogramming and ES/EB differentiation

The ATAC-seq dataset was downloaded from NCBI GEO database (GSE168636)^35^. Sequence reads for undifferentiated MEFs were pooled from two replicates (GSM5151877 and GSM5151879), and for Flk1^+^ MEFs at day 7 post-Etv2 induction were pooled from two replicates (GSM5151861 and GSM5151863). Sequence reads for undifferentiated EBs were pooled from two replicates (GSM5151873 and GSM5151875), and for Flk1^+^ EBs at day 2.5 post Etv2 induction were pooled from two replicates (GSM5151869 and GSM5151871). MACS2 (v2.1.1) identified 57,732 peaks for undifferentiated MEFs and Flk1+ MEFs at day 7 post Etv2 induction and 36,114 peaks for undifferentiated EBs and Flk1+ EBs at day 2.5 post Etv2 induction. We used motifmatchr (v1.16.0) to obtain 20,822 and 24,935 putative Etv2 motif binding regions for MEFs and EBs, respectively,

#### Ascl1 induced MEF reprogramming

The ATAC-seq dataset was downloaded from NCBI GEO database (GSE101397)^25^. The sequence reads for undifferentiated MEFs was obtained from GSM2701947. For MEFs at day 22 post Ascl1 induction the sequence reads were pooled from three replicates (GSM2701979, GSM2701980 and GSM2701981). MACS2 identified 123,271 peaks for undifferentiated MEFs and MEFs at day 22 post Ascl1 induction and motifmatchr identified 71,616 canonical Ascl1 motif binding sites.

#### OSK induced MEF reprogramming

The ATAC-seq dataset was downloaded from NCBI GEO database (GSE93026)^47^. The sequence reads for the undifferentiated MEF samples were pooled from two replicates (GSM2442671 and GSM2442671), and for the MEFs at day 7 post-OSK induction were pooled from two replicates (GSM2442705 and GSM2442706). We used motifmatchr to identify 282,789 putative binding sites for Oct4, Sox2 or Klf4 for the downstream analysis.

#### ATAC-seq preprocessing

The sequencing reads where mapped to the mouse and human genome (mm10 or hg19) using Bowtie2 (v2.2.4)^62^. The ATAC-seq reads lied on chromosome Y and mitochondria were excluded^63^. ChromVAR (v1.10)^51^ were used for transcription factor based chromatin accessibility analysis. 322 transcription factors compiled in the Homer database were used for the chromVAR analysis. The pathway analysis was performed using R packages clusterProfiler and ChIPseeker^64,65^.

## Code availability

SeATAC is available at https://github.com/gongx030/seatac as an R package. Codes pertaining to important analyses in this study are available from GitHub webpage (https://github.com/gongx030/seatac_manuscript).

