## Supplementary Figures for "SeATAC: a tool for exploring the chromatin landscape and the role of pioneer factors"

### Supplementary Figure 1

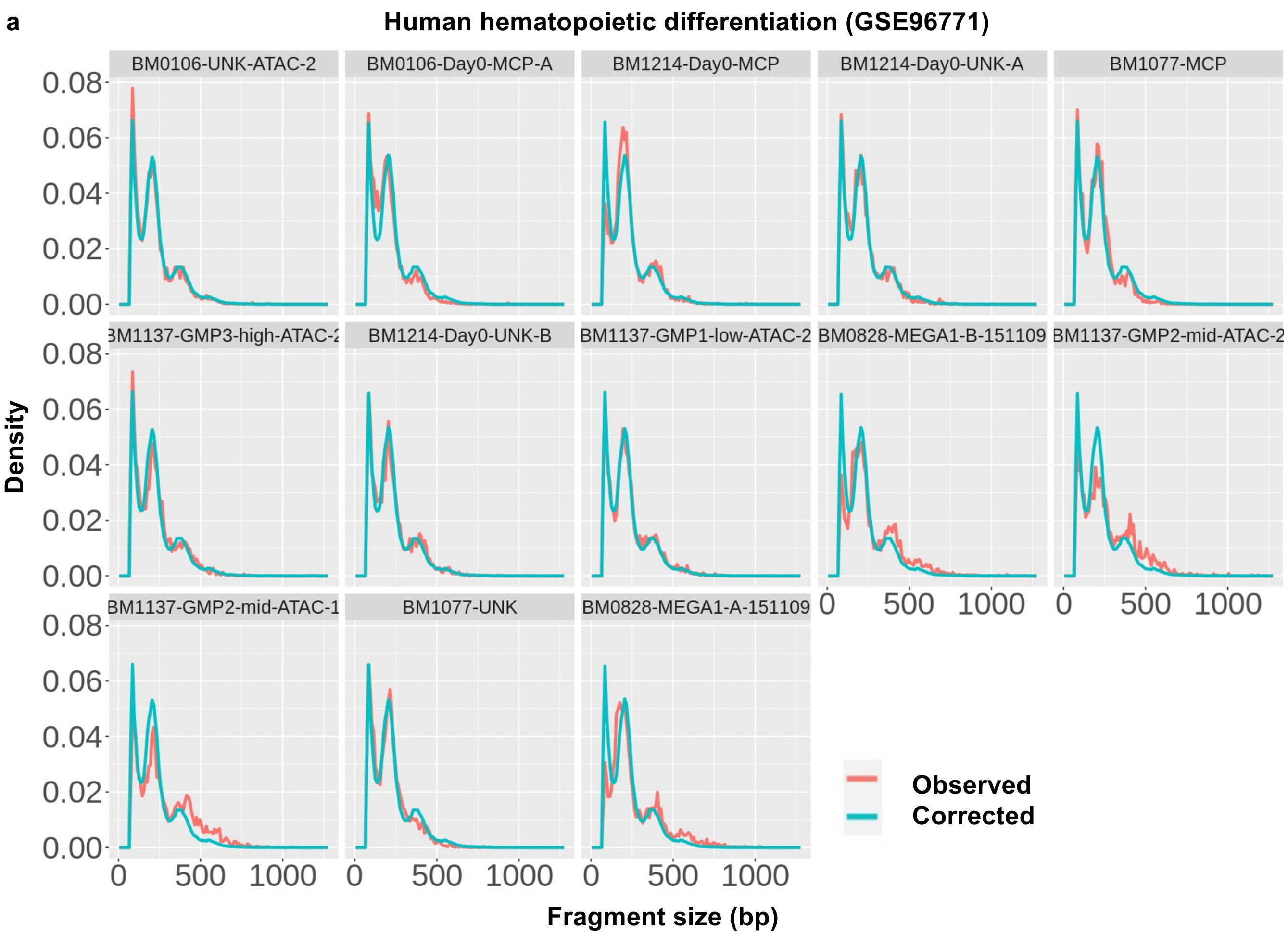

**Supplementary Figure 1. SeATAC corrects batch effects in fragment size distributions in ATAC-seq data. (a)** The density plots show the observed (red) and corrected (green) fragment size distribution of 13 samples from a human hematopoietic differentiation ATAC-seq data (GSE96771). The x-axis shows the fragment size in base pairs (bp) and the y-axis shows the density.

### Supplementary Figure 2

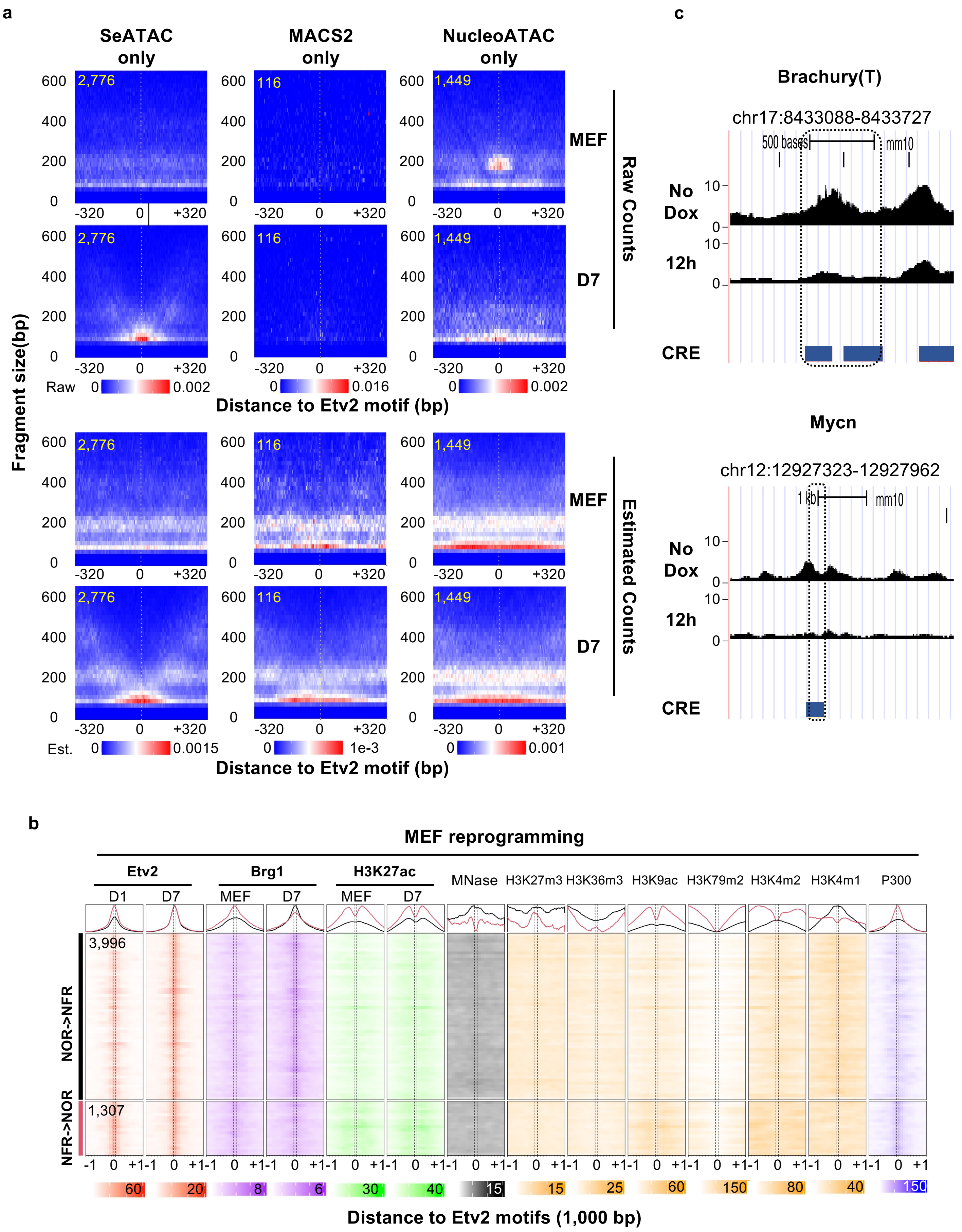

**Supplementary Figure 2. SeATAC detects Etv2 binding sites with increased chromatin accessibility during Etv2 induced EB differentiation and MEF reprogramming. (a)** The aggregated V-plot includes 2,776, 116 and 1,449 Etv2 motifs with increased chromatin accessibility identified by SeATAC only, MACS2 only and NucleoATAC only using the ATAC-seq dataset of Etv2 induced MEF reprogramming (undifferentiated MEFs vs. Flk1$^+$ cells at 7 days post-induction). Both raw V-plots and estimated V-plots are shown. The heatmap color indicates the normalized read density for raw counts (top) and the estimated read density for estimated read counts (bottom). **(b)** The heatmaps show the Etv2, Brg1, H3K27ac ChIP-seq of 3,996 and 1,307 Etv2 binding sites that have increased (NOR->NFR) or decreased (NFR->NOR) chromatin accessibility in undifferentiated MEFs (Brg1 and H3K27ac), 1 day post-Etv2 induction (Etv2), and 7 days post-Etv2 induction (Etv2, Brg1 and H3K27ac). The heatmaps also include the MNase-seq, H3K27m3, H3K36m3, H3K9ac, H3K79me2, H3K4me2, H3K4me1 and P300 ChIP-seq from undifferentiated MEFs. **(c)** The UCSC genome browser track show the ATAC-seq density near the Etv2 motifs at the promoters of Brachyury (T) and Mycn. The Etv2 motifs become less accessible during the EB differentiation and both Brachyury (T) and Mycn are significantly down-regulated during Etv2 induced EB differentiation.

### Supplementary Figure 3

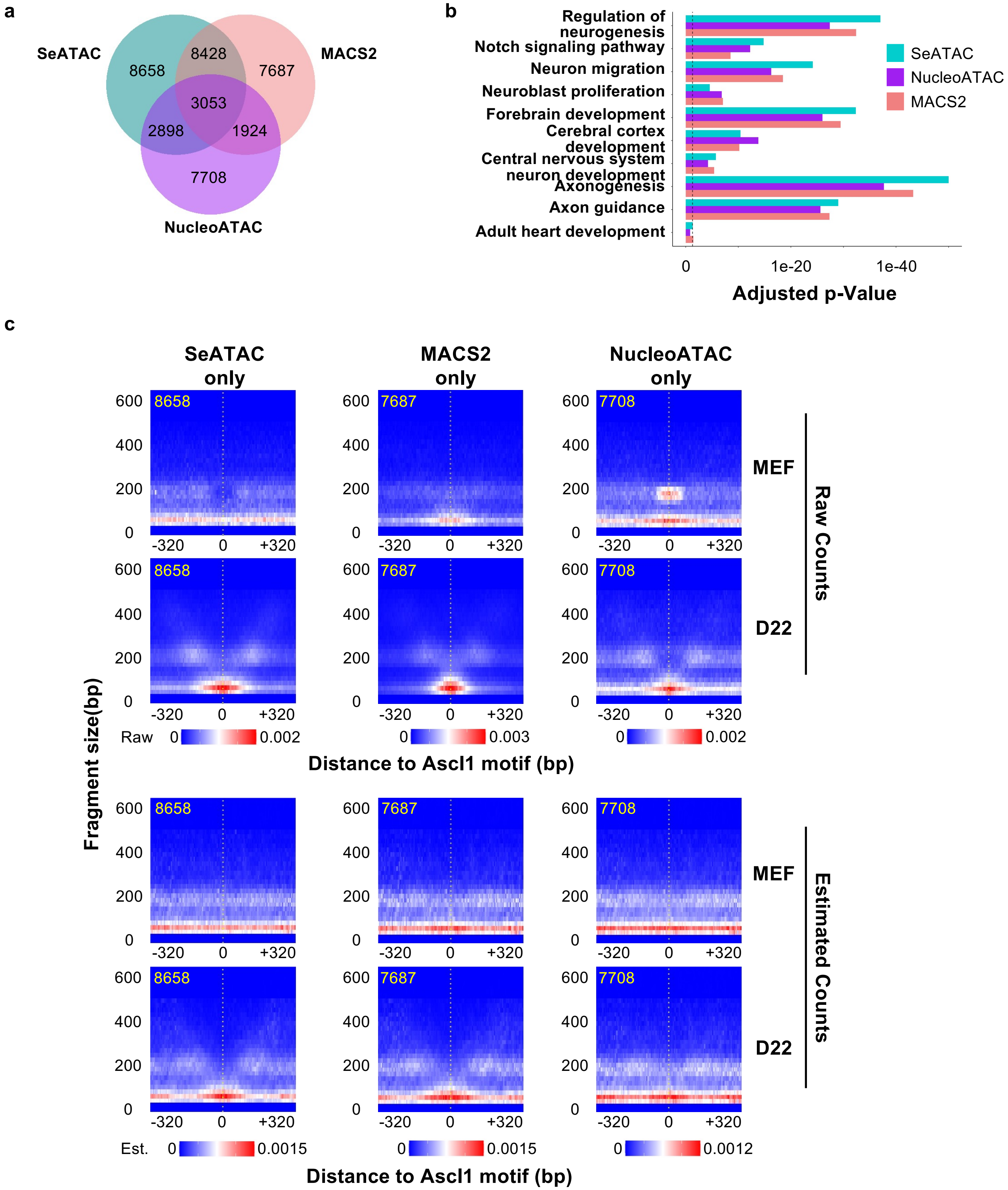

**Supplementary Figure 3. SeATAC detects Ascl1 binding sites with increased chromatin accessibility during Ascl1 induced MEF reprograming. (a)** The Venn diagrams show the number of Ascl1 motifs with increased chromatin accessibility identified using SeATAC, MACS2 and NucleoATAC. **(b)**The barplots show the Gene Ontology (GO) terms that are significantly associated with the genes that have promoters (-5,000 - +1,000bp region flanking the TSS) with Ascl1 binding motifs with increased chromatin accessibility, identified by SeATAC, MACS2 and NucleoATAC. The y-axis showed the adjusted *p*-value of the pathway analysis. **(c)** The aggregated V-plot includes 8,658, 7,687 and 7,708 Ascl1 motifs with increased chromatin accessibility identified using SeATAC only, MACS2 only and NucleoATAC only in ATAC-seq data of Ascl1 induced MEF reprogramming (undifferentiated MEFs vs. 22 days post Ascl1 induction). Both raw V-plots and estimated V-plots are shown. The heatmap color indicates the normalized read density for raw counts (top) and the estimated read density for estimated read counts (bottom).

### Supplementary Figure 4

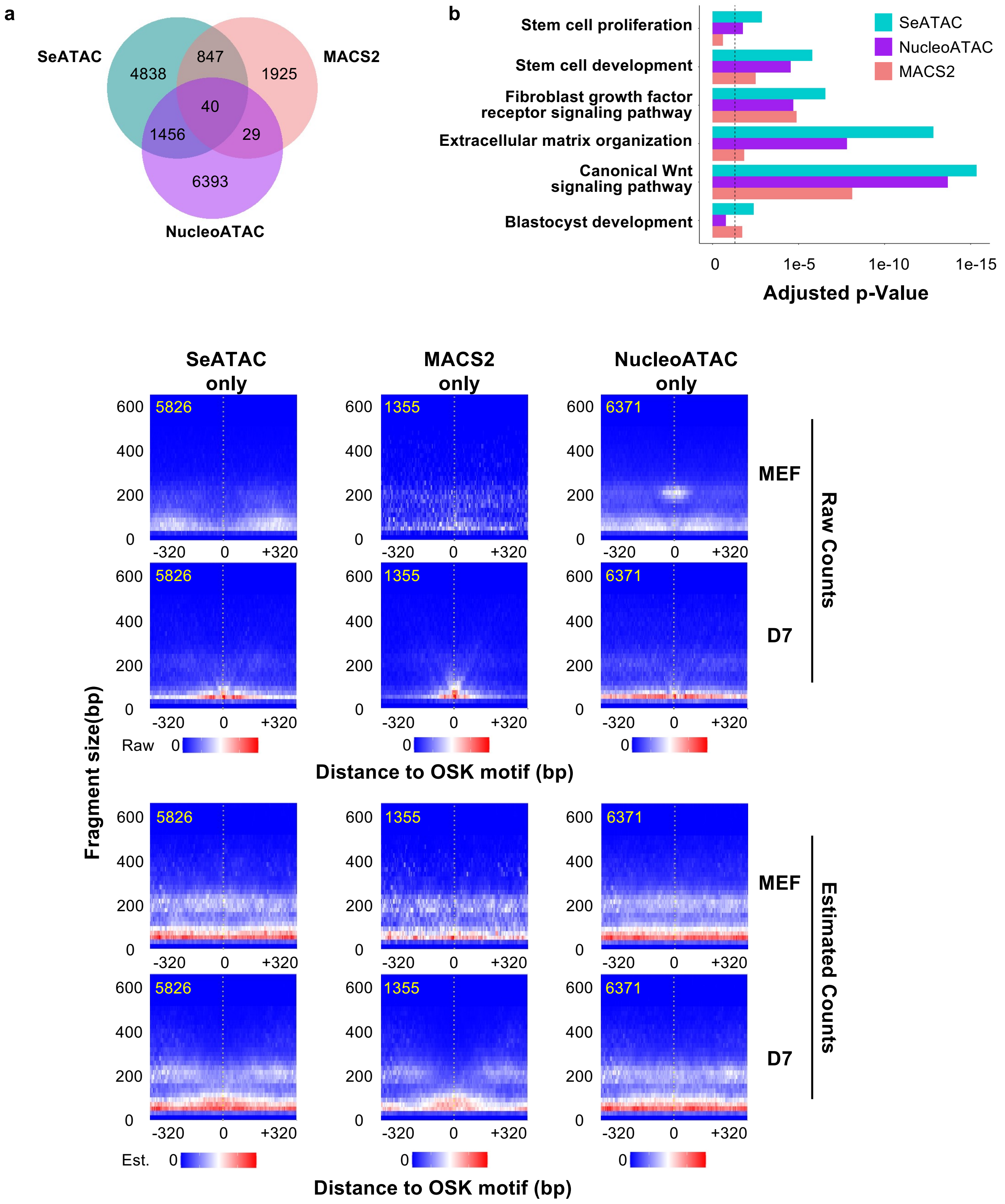

**Supplementary Figure 4. SeATAC detects OSK binding sites with increased chromatin accessibility during OSK induced reprograming. (a)** The Venn diagrams show the number of OSK motifs with increased chromatin accessibility identified using SeATAC, MACS2 and NucleoATAC. **(b)**The barplots show the Gene Ontology (GO) terms that are significantly associated with the genes with promoters (-5,000 - +1,000bp region flanking the TSS) that have OSK motifs with increased chromatin accessibility, identified by SeATAC, MACS2 and NucleoATAC. The y-axis showed the adjusted *p*-value of the pathway analysis. **(c)** The aggregated V-plot includes 5,826, 1,355 and 6,371 OSK motifs with increased chromatin accessibility identified using SeATAC only, MACS2 only and NucleoATAC only in ATAC-seq data of OSK induced MEF reprogramming (undifferentiated MEFs vs. 7 days post OSK induction). Both raw V-plots and estimated V-plots are shown. The heatmap color indicates the normalized read density for raw counts (top) and the estimated read density for estimated read counts (bottom).

### Supplementary Figure 5

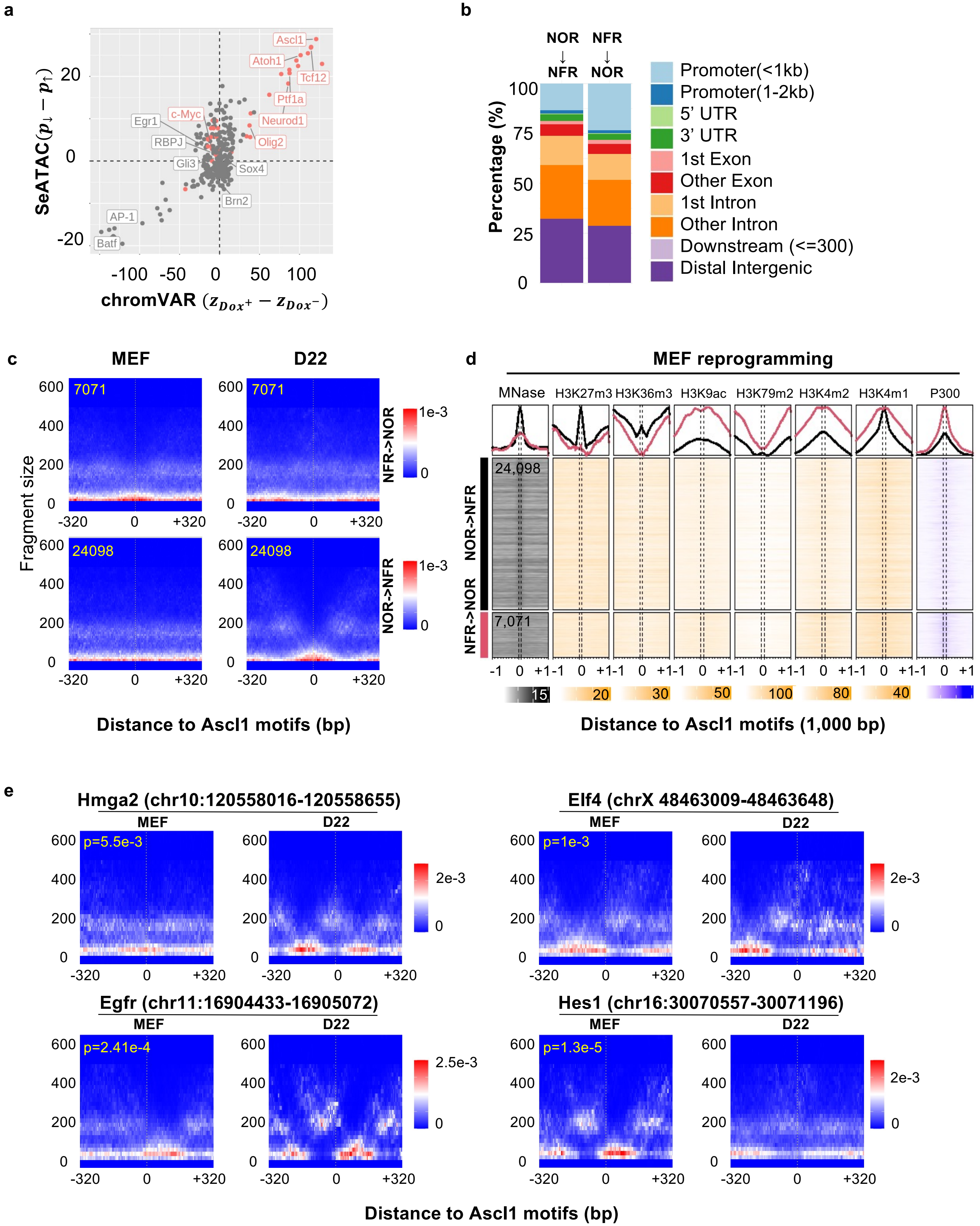

**Supplementary Figure 5. Induction of Ascl1 causes both chromatin relaxation and closure at Ascl1 binding sites. (a)** The dot plots compare the changes of motif associated chromatin accessibility estimated by chromVAR (x-axis) and the difference of the percent of TFBS with decreased or increased chromatin accessibility estimated by SeATAC (y-axis). $\boldsymbol{z}_{\boldsymbol{Dox}^+}$ and $\boldsymbol{z}_{\boldsymbol{Dox}^-}$ are the normalized deviation score of $Dox^+$ condition (22 days post-induction for MEF reprogramming) and $Dox^-$ condition (undifferentiated MEFs). $\boldsymbol{p}_{\downarrow}$ and $\boldsymbol{p}_{\uparrow}$ are the percent of TFBS that shows decreased or increased chromatin accessibility in $Dox^+$ condition compared with the $Dox^-$ condition. **(b)** The barplots show the genomic distribution of Ascl1 binding sites with decreased (NFR->NOR) or increased (NOR->NFR) chromatin accessibility in MEF reprogramming. The change of chromatin accessibility is estimated using SeATAC. **(c)** The aggregated V-plot includes: 24,098 and 7,071 Ascl1 binding sites that have increased (NOR->NFR) or decreased (NFR->NOR) chromatin accessibility during MEF reprograming. The heatmap color indicates the estimated read density. **(d)** The heatmaps show the MNase-seq, H3K27m3, H3K36m3, H3K9ac, H3K79me2, H3K4me2, H3K4me1 and P300 ChIP-seq signals in undifferentiated MEFs of 24,098 and 7,071 Ascl1 binding sites that have increased (NOR->NFR) or decreased (NFR->NOR) chromatin accessibility during the MEF reprogramming. The change of chromatin accessibility is estimated using SeATAC. **(e)** The V-plot show Ascl1 motifs with decreased chromatin accessibility at the promoters (-5,000 - +1,000bp region flanking the TSS) of four genes (Hmga2, Elf4, Egfr and Hes1) that are down-regulated during the Ascl1 induced MEF reprogramming.

### Supplementary Figure 6

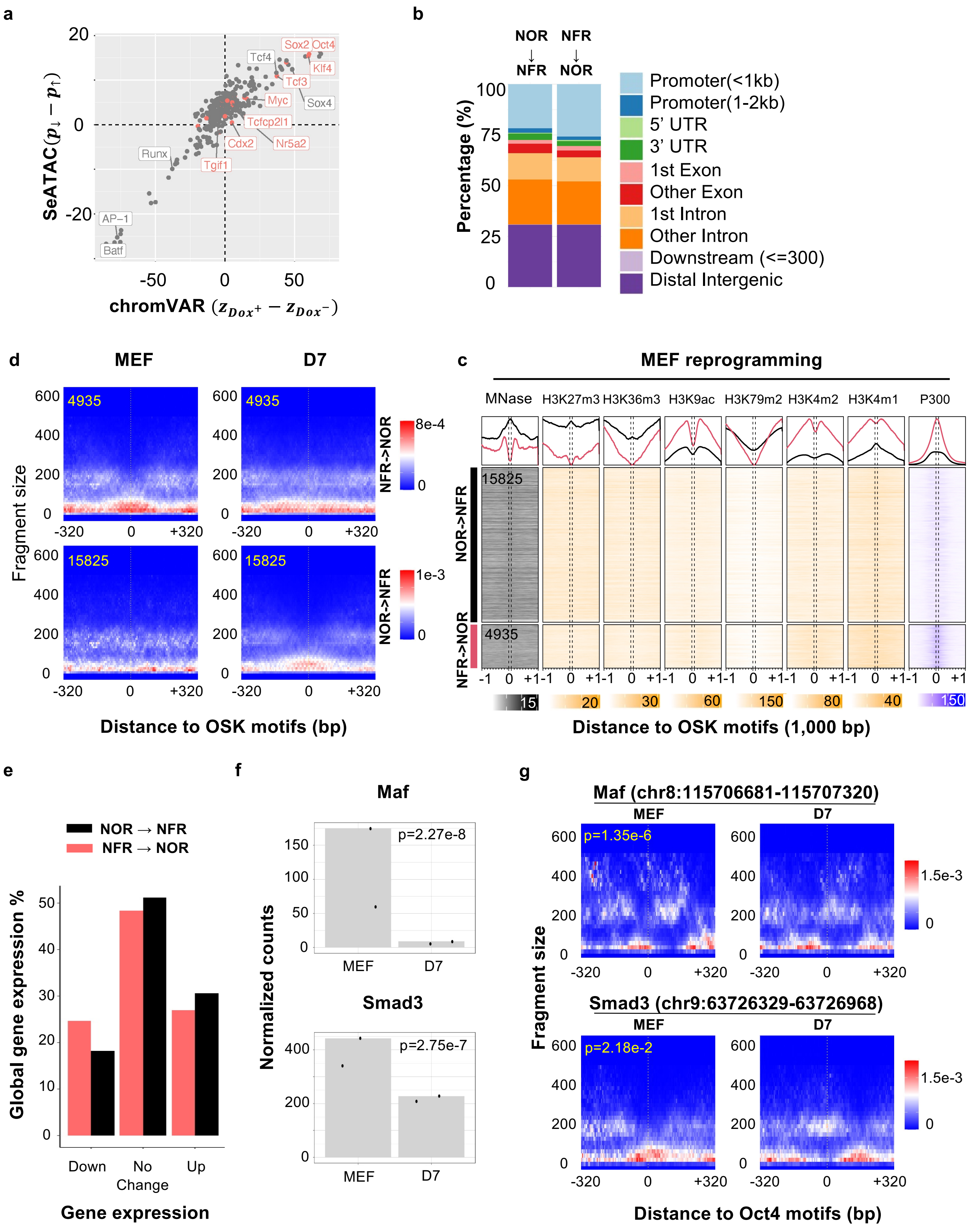

**Supplementary Figure 6. Induction of OSK (Oct4/Sox2/Klf4) causes both chromatin relaxation and closure at OSK binding sites. (a)** The dot plots compare the changes of motif associated chromatin accessibility estimated by chromVAR (x-axis) and the difference of the percent of TFBS with decreased or increased chromatin accessibility estimated by SeATAC (y-axis). $\boldsymbol{z_{Dox^+}}$ and $\boldsymbol{z_{Dox^-}}$ are the normalized deviation score of $Dox^+$ condition (7 days post-induction for MEF reprogramming) and $Dox^-$ condition (undifferentiated MEFs). $\boldsymbol{p_{\downarrow}}$ and $\boldsymbol{p_{\uparrow}}$ are the percent of TFBS that shows decreased or increased chromatin accessibility in $Dox^+$ condition compared with the $Dox^-$ condition. **(b)** The barplots show the genomic distribution of OSK binding sites with decreased (NFR->NOR) or increased (NOR->NFR) chromatin accessibility in MEF reprogramming. The change of chromatin accessibility is estimated using SeATAC. **(c)** The aggregated V-plot include 15,825 and 4,935 OSK binding sites that have increased (NOR->NFR) or decreased (NFR->NOR) chromatin accessibility during MEF reprograming. The heatmap color indicates the estimated read density. **(d)** The heatmaps show the MNase-seq, H3K27m3, H3K36m3, H3K9ac, H3K79me2, H3K4me2, H3K4me1 and P300 ChIP-seq signals in undifferentiated MEFs of 15,825 and 4,935 OSK binding sites that have increased (NOR->NFR) or decreased (NFR->NOR) chromatin accessibility during the MEF reprogramming. The change of chromatin accessibility is estimated using SeATAC. **(e)** The barplots show the percent of genes that were down-regulated, up-regulated or not changed between undifferentiated MEFs and 7 hours post OSK induction. **(f-g)** Maf and Smad3 **(f)** are significantly down-regulated during the OSK induced MEF reprogramming and **(g)** have OSK motifs that become significantly less accessible during the differentiation at their promoter region (-5,000 - +1,000bp region flanking the TSS). The heatmap color indicates estimated read density.
